# Reconstruction of epithelial transcriptional trajectories reveals heterogeneous progression and early therapy-resistance programs in high-grade serous carcinoma precursors

**DOI:** 10.64898/2026.02.06.704093

**Authors:** Michail Sideris, Eleni Maniati, Tanjina Kader, Sandro Santagata, Ronny Drapkin, Frances Balkwill, Ranjit Manchanda

**Author notes:** Corresponding author **Corresponding author:** Ranjit Manchanda, MD, PhD, Centre for Cancer Screening, Prevention & Early Diagnosis (CCSPED), Wolfson Institute of Population Health, Queen Mary University of London, Charterhouse Square, EC1M 6BQ, London, UK. Equal Contribution (joint first) (Michail Sideris, Eleni Maniati). Equal Contribution (Frances Balkwill, Ranjit Manchanda).

## Abstract

**Background:** Serous-tubal intraepithelial-carcinoma (STIC) is considered the principal precursor of tubo-ovarian high-grade serous-carcinoma (HGSC), yet its biological uniformity, progression risk and potential response to poly(ADP-ribose) polymerase-inhibitors (PARPi) remain poorly defined.

**Methods:** We performed trajectory-based transcriptomic reconstruction of Fallopian-tube epithelial regions spanning normal epithelium, precursor lesions, STIC, and invasive carcinoma using pseudotime inference and publicly available spatial transcriptomic data. Gene expression and pathway dynamics were defined along pseudotime, and interpatient heterogeneity examined at both lesion and patient levels. Transcriptional signatures associated with PARPi-resistance were quantified across STICs, *BRCA*-mutant and *BRCA*-wild type subtypes.

**Results:** Trajectory inference captured a continuous transcriptional progression from normal epithelium to HGSC, with STICs occupying heterogeneous evolutionary positions rather than a single precursor state. Incidental isolated STICs(STICi) spanned early to later pseudotime states and frequently aligned with loss-of-cilium organisation and less advanced epithelial phenotypes. STICs associated with concurrent cancer(STICc) exhibited more advanced malignant progression signatures, including cell-cycle activation, epithelial-to-mesenchymal transition (EMT), interferon signaling, and DNA-repair. Histologically similar lesions occupied divergent pseudotime positions with marked interpatient heterogeneity. PARPi resistance–associated signatures were variably enriched across precursor and precancer-stage lesions, with persistence into invasive disease.

**Conclusions:** STICs are heterogeneous and occupy distinct evolutionary positions along a continuum, highlighting potentially different progression risks from normal epithelium to HGSC. STICi differ from STICc which harbour signatures of more advanced malignant progression. Heterogeneity in PARPi resistance–associated programs in STICs cautions against uniform (non-stratified) use of PARPi-based primary prevention strategies. Future research should explore evolutionary-trajectory informed biomarkers for risk stratification and early interception strategies.

**Translational relevance:** Serous tubal intraepithelial-carcinoma (STIC) represents a precursor of tubo-ovarian high-grade serous carcinoma (HGSC), offering unique translational opportunities for early detection, risk stratification, and prevention. However, its phenotypic uniformity, progression risk and therapeutic drug response are not fully understood. Our study shows that STICs are not a uniform precursor state, but instead occupy a range of evolutionary positions along a continuum from normal epithelium to invasive HGSC. This heterogeneity reflects continuous transcriptional variation rather than discrete precursor categories. We report gene expression dynamics and molecular signature changes across the malignant transformation timeline. We further illustrate that precursor lesions exhibit phenotypic variability that impacts therapeutic drug resistance-associated programs. Our data highlight potential disease biomarkers defined by evolutionary trajectory inference and transcriptional processes associated with drug resistance operating in precursor lesions. These findings may help improve our ability to distinguish clinically significant lesions and inform targeted interception strategies.

## Introduction

Serous tubal intraepithelial carcinoma (STIC) is an established precursor of tubo-ovarian high-grade serous carcinoma (HGSC).(1, 2) Evolutionary genomic analyses support a clonal continuum in which malignant progression initiates in the Fallopian tube(1, 3) and indicate an estimated window of 6-7 years between development of a STIC and initiation of HGSC.(1, 4) Nevertheless, our understanding of the biological processes underlying early disease evolution remains limited. Isolated incidental STICs (STICi) may represent biologically indolent lesions or more advanced precursors comparable to STICs associated with concurrent cancer (STICc). Whether these lesions occupt distinct of overlapping evolutionary states remains unclear. These distinctions have direct implications for progression risk and therapeutic management, including sensitivity to poly(ADP-ribose) polymerase inhibitors (PARPi). Data from HGSC cell lines implicate transcriptional programs associated with attenuated replication stress as markers of PARPi resistance.(5)

The recently published multimodal atlas of Fallopian tube precursors and HGSC(6) encompasses spatial transcriptomic profiling of 407 epithelial regions of interest (ROIs), including normal epithelium (Fallopian tube and/or fimbriae), precancer lesions, and regions of invasive cancer (HGSC), derived from 34 patients. We applied trajectory inference to this dataset to reconstruct the disease progression continuum, define transcriptomic changes across pseudotime, and characterise transcriptional heterogeneity among Fallopian tube precursor lesions. Finally, we quantified enrichment of a PARPi resistance-associated transcriptional signaturse in STICs and related these to evolutionary position and malignant transformation phenoptypes.

## Methods

### Dataset features

The Kader et al. (6) GeoMx dataset, containing normalized gene expression values and clinical sample metadata, was obtained from the GEO repository (GSE281193). The dataset contained two groups of specimens: Group 1 encompassing specimens with HGSC and co-occurring STICs (hereforth referred to as “STICc”); and Group 2 lacking HGSC but containing isolated precursor lesions identified incidentally during risk-reducing salpingo-oophorectomy (RRSO) or opportunistic salpingectomy (hereforth referred to as “STICi”). ROI annotations followed those provided in the Kader et al. dataset (see Supplementary Table-S1). Twenty-five of the 34 specimens with epithelial ROIs had matched “normal” Fallopian tube and/or fimbriae epithelium within the same tissue section. Serous tubal intraepithelial lesions (STIL) were included. Cancer regions detached from the surrounding tissue were denoted as “floating cancer”. The number of ROIs per lesion type per group, along with relevant abbreviations, is listed in Supplementary Table-S1.

### Trajectory inference using slingshot

Pseudotemporal ordering of the epithelial ROIs was performed using Slingshot.(7) Slingshot integrates clustering information with dimensionality reduction to infer lineage relationships using a minimum spanning tree and subsequently fits smooth principal curves to model continuous transcriptional progressions. Prior to trajectory inference, the 2,000 most variable genes in the dataset were selected. Principal component analysis (PCA) was performed on the normalized expression matrix, and the first four principal components were used as input for Slingshot. Trajectory inference was conducted using the lesion-type annotations as cluster labels to guide lineage construction. The analysis was performed in an unsupervised manner without pre-specifying start or end clusters, allowing slingshot to infer the trajectory structure and directionality based on the data topology. Slingshot constructed minimum spanning trees connecting clusters and fitted principal curves to define smooth trajectories. Pseudotime values were calculated for each ROI along each identified lineage, representing relative transcriptional position along the trajectory. ROIs not assigned to a given lineage were assigned NA pseudotime values for that lineage.

### Gene set variation analysis

Gene set variation analysis (GSVA) was conducted using the GSVA package, and heatmaps were generated using ComplexHeatmap. GSVA derives enrichment scores for each sample as a function of genes inside and outside a given gene set, analogous to a competitive gene set test.(8) This approach converts a gene–by-sample matrix, providing an estimate of pathway activity.(8) Gene Ontology Biological Processes (GOBP) and Hallmark gene sets were obtained from the human MSigDB collections. A transcriptional signature associated with PARPi resistance, defined by Tamura et al. (5) and focusing on upregulated genes (Supplementary Table-S6), was used to evaluate PARPi resistance-associated programs in the Kader dataset. We first assessed the performance of this signature in ovarian cancer cell lines from the Cancer Cell Line Encyclopedia (CCLE) dataset (9), confirming a significant positive correlation with olaparib IC50 values (mean of two experimental replicates). We then evaluated associations between this signature and GOBP GSVA scores in STIC ROIs and invasive HGSC using Pearson’s correlation, with Benjamini-Hochberg correction for multiple testing. Additional transcriptional signatures related to attenuated replication stress, paclitaxel resistance, and congression defects (defined in Tamura et al.) (5) were included to further examine processes linked to chromosomal instability (Supplementary Table S6).

Word clouds were generated using tidytext and wordcloud. All analyses were performed in R version 4.4.2.

## Results

Trajectory analysis provides a valuable framework to investigate disease progression from transcriptomic data and capture gradual transitions underlying pathological processes, particularly in contexts where longitudinal sampling or explicit time-course measurements are infeasible.(7) Here, we applied trajectory inference using Slingshot(7) on GeoMx transcriptomes from 407 epithelial ROIs spanning Fallopian tube precursor lesions, invasive cancer, and normal tissue in the Kader dataset,(6) corresponding to 34 patients, to reconstruct HGSC progression. In principle, this approach estimates relative transcriptional ordering based on similarities in gene exprssion profiles, enabling assignment of pseudotime values to individual ROIs. Importantly, pseudotime reflects relative evolutionary position rather than chronological time.

Trajectory inference reconstructed a continuous transcriptional evolution from normal Fallopian tube epithelium to HGSC. Principal component embedding revealed a smooth disease trajectory across epithelial ROIs (Figure 1A, Supplementary Figure S1A-B). Pseudotime increased stepwise across lesion types, from normal Fallopian tube/fimbriae and p53 signatures lesions through STIL, STICi, STICc, and ultimately invasive and floating cancer, capturing known biological and histological phenotypes (Figure 1B, Supplementary Table-S1, Supplementary Table-S2). We observed a statistically significant difference in pseudotime distribution among p53 signature lesions and invasive cancer cells in *BRCA*mut vs *BRCA*wt cases (Figure S1C), consistent with the earlier clinical stage typically observed in *BRCA*mut specimens.

**Figure 1.**
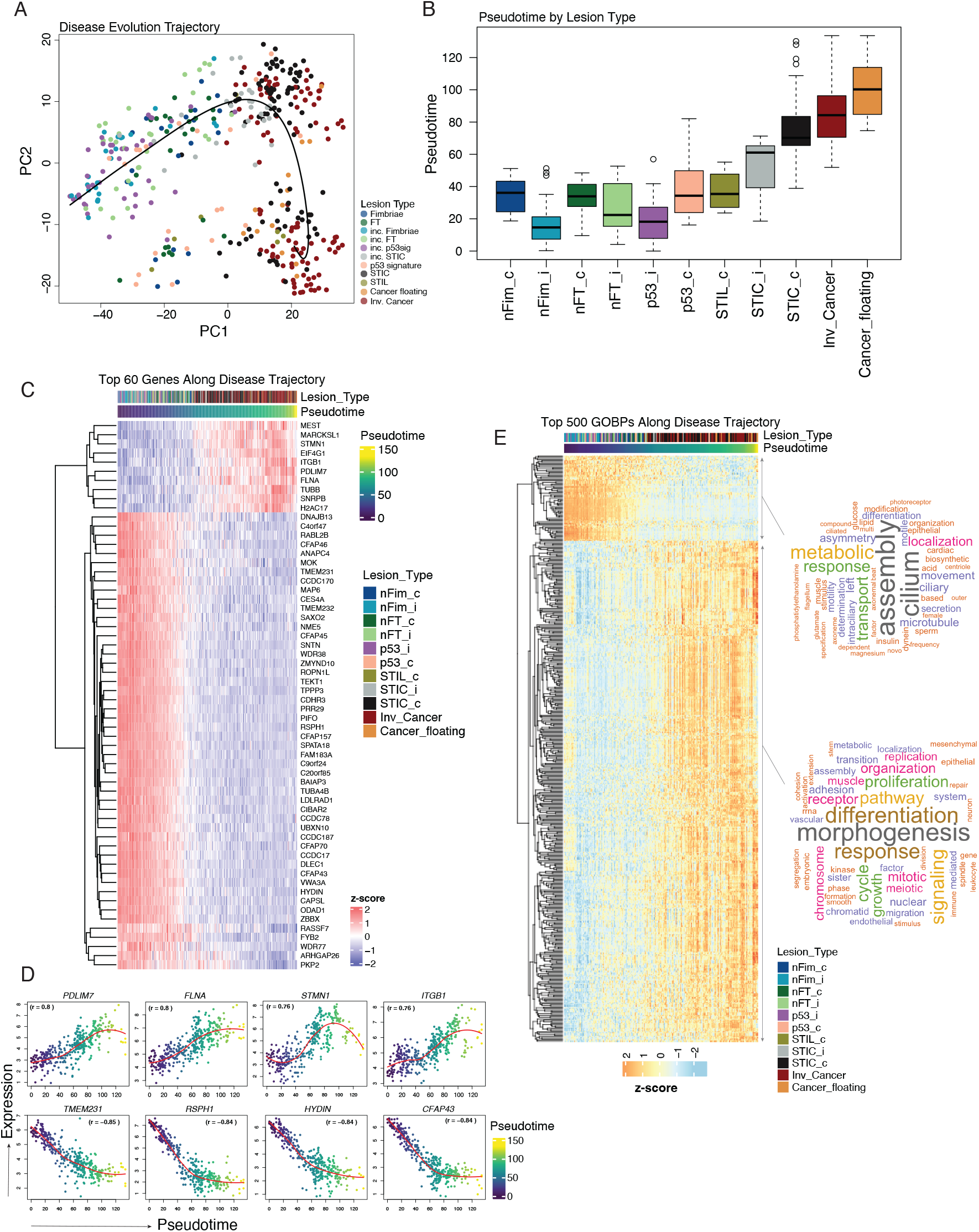
Pseudotemporal reconstruction of transcriptional changes from normal to invasive ovarian cancer epithelium. Principal Component Analysis (PCA) of the Kader dataset using the top 2,000 most variable genes, each dot is an epithelial region of interest (ROI, n = 407 ROIs), coloured by lesion type. **B)** Boxplot illustrating ROIs pseudotime values for each lesion type (Kruskal-Wallis p = 8.68e-56). **C)** Heatmap of normalised expression of the top sixty genes that significantly change in expression with pseudotime along the first slingshot trajectory of the Kader et al. dataset (Spearman correlation with Benjamini-Hochberg, BH, adjusted p < 0.05). **D)** Scatterplots illustrating top correlating genes (*PDLIM7, FLNA, STMN1, ITGB1, TMEM231, RSPH1, HYDIN, CFAP43*). Each dot corresponds to an ROI, ordered and coloured by pseudotime. **E)** Heatmap of gsva enrichment scores for top five hundred Gene Ontology Biological Processes (GOBPs) that significantly change in expression with pseudotime, along the first slingshot trajectory (Spearman BH adjp < 0.05, r > 0.5). Word cloud graph highlights common keywords of the up and down-regulated processes.

Coordinated gene expression remodeling along pseudotime was evident (Figure 1C; Supplementary Table S3), with two distinct clusters of genes progressively downregulated or upregulated along the evolutionary trajectory to HGSC. Top genes positively correlated with pseudotime included the cytoskeletal organisation genes *PDLIM7* and *FLNA*, the microtubule regulator *STMN1* – a known immunohistochemical marker of STIC lesions(10) – and *ITGB1*, which is involved in cell adhesion and migration (Figure 1D, Supplementary Table-S3). By contrast, genes negatively correlated with pseudotime included *TMEM231, RSPH1, HYDIN*, and *CFAP43*, all of which are related to ciliary structure and function (Figure 1D, Supplementary Table-S3). Each dot of the scatterplots corresponds to a ROI, demonstrating a continuum in expression trend across the pseudotime.

Pathway level analysis demonstrated progressive loss of cilium organisation and upregulation of embryonic morphogenesis, proliferation, chromosome segregation, adhesion and epithelial-to-mesenchymal transition (EMT) (Figure 1E, Supplementary Table-S4). We further observed upregulation of TGF-β signaling accompanied by IL6/STAT3, MYC and TNF-α signalling via NF-κB (Figure 1E, Figure S1D, Supplementary Table-S5). Together, these changes indicate malignant transformation occurring alongside increasing pseudotime through coordinated disruption of epithelial structure and activation of oncogenic and inflammatory transcriptional programmes well known for promoting cancer growth and spread.(11-13)

We next created a pseudotime position of each lesion at the patient level (Figure 2A) to assess interpatient heterogeneity cross the malignant transformation continuum. This analysis revealed substantial variability among lesions of the same histologic type. For example, STICi lesions from patient 43 occupied pseudotime positions similar to STICc lesions in patients 6 and 25, yet were positioned later than STICi lesions in patient 36.

**Figure 2.**
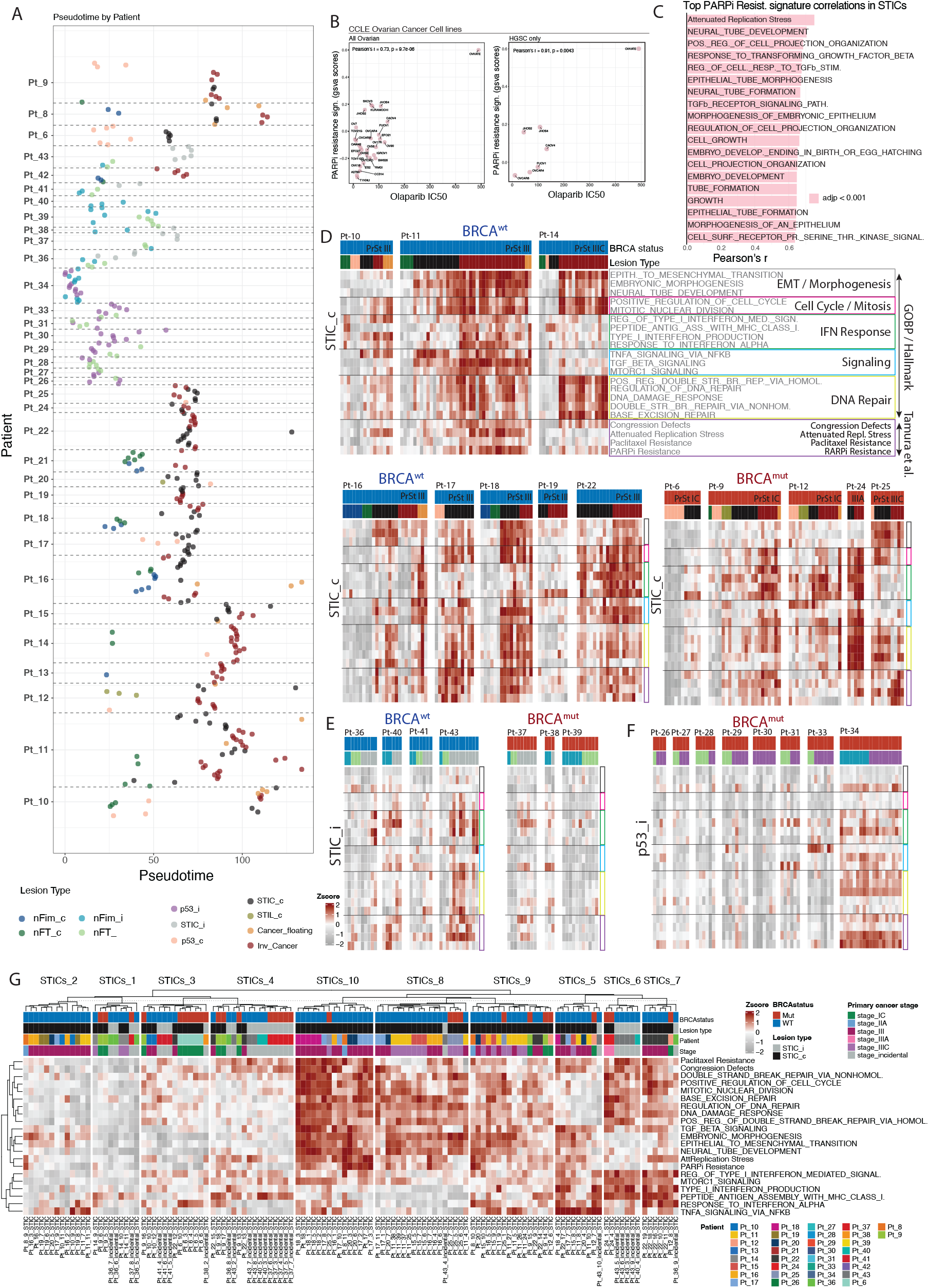
“Chronotope” of lesions per patient and transcriptional features of therapy resistance. **A)** Scatter plot illustrating the position of each lesion in pseudotime, per patient. Colours correspond to lesion type. **B)** Scatter plot illustrating correlation of PARPi resistance signature (Tamura et al) with Olaparib IC50 in ovarian cancer cell lines in the CCLE dataset (n = 28 ovarian cancer cell lines, n = 7 HGSC cells lines). **C)** Barplot of STIC ROI transcriptomes showing top 10 Biological Processes and transcriptional signatures related to Congression Defects, Attenuated Replication Stress, Paclitaxel Resistance significantly correlating with a PARP-inhibitor resistance signature from Tamura et al (pearson’s BH adjp < 0.05). **D–F)** Heatmaps of significant disease related GOBPs and Hallmarks as well as the Tamura et al transcriptional signatures for **D)** patients with invasive cancer **E)** patients with incidental STIC lesions and **F)** patients with incidental p53 lesions. Terms have been grouped in the following categories: EMT / morphogenesis (Epithelial to Mesenchymal Transition, Embryonic Morphogenesis, Neural Tube Development); cell cycle / mitosis (Positive Regulation of Cell Cycle, Mitotic Nuclear Division); IFN response (Regulation of Type I Interferon–Mediated Signaling Pathway, Peptide Antigen Assembly with MHC Class I Protein Complex, Type I Interferon Production, Response to Interferon-Alpha); Oncogenic Signaling ( TNFA signaling via NFkB, TGF-beta signaling, MTORC1 signaling); DNA repair (Positive Regulation of Double-Strand Break Repair via Homologous Recombination, Regulation of DNA Repair, DNA Damage Response, Double-Strand Break Repair via Nonhomologous End Joining, Base Excision Repair) as well as transcriptional signatures from Tamura et al. D-F) Lesion type colour coding as in A. **G)** Heatmap of gsva enrichment scores showing clustering patterns of STIC ROIs using the processes and pathways described in the previous panel (D-F). Clustering of ROIs was performed by k-means clustering.

Enrichment of a published PARPi resistance-associated transcriptional signature correlated with increasing olaparib IC50 values of ovarian cancer cell lines from the Cancer Cell Line Encyclopedia (CCLE) dataset (Figure 2B) (5, 9). Within STICs, this signature was strongly associated with pathways linked to attenuated replication stress, neural tube development, TGF-β response, and embryonic morphogenesis (Figure 2C, Supplementary Table-S6 and Supplementary Table-S7). We further observed significant positive associations with WNT signaling and EMT (adjusted p < 0.0001, r > 0.6; Supplementary Table-S6). Similar associations were observed in HGSC; however, the relative ranking differed, with embryonic morphogenesis, WNT signaling, and TGF-β response emerging as the most strongly correlated processes (Figure S2A, Supplementary Table-S8).

Patient-level pathway analyses (Figures 2D–F and S2B) revealed heterogeneous transcriptional profiles across EMT/morphogenesis, cell-cycle progression, interferon response, oncogenic signaling, DNA repair, congression delays, replication stress attenuation, and paclitaxel and PARPi resistance programs. In patients with concurrent HGSC (Figure 2D), some STICc lesions exhibited elevated EMT, proliferation, and DNA repair activity (e.g., patients 11, 17, 18, and 25), features that were largely preserved in matched invasive HGSC regions. By contrast, certain cases showed discordant DNA repair activity between STICc and invasive HGSC (e.g., patients 14, 16, and 25). PARPi resistance-associated activity was observed in a large number of STICc lesions (patients 11, 16, 17, 18, and 22), occassioanlly co-occurring with paclitaxel resistance. Notably, patient 18 displayed elevated PARPi resistance-associated programs in STICs but reduced activity in matched invasive regions.

Several patients displayed strong interferon responses in HGSC (patients 9, 11, 12, 22, and 24), with corresponding STICc lesions showing variable interferon activity ranging from low (patient 11) to similarly elevated (patient 22 and 24). STICi lesions generally showed low EMT and proliferation activity in most patients, in keeping with their low DNA repair activity (Figure 2E; patients 36–41). These resistance-associated signatures had variable activity in STICi, albeit at lower levels compared to STICc (Figure 2E).

Of particular interest, patient 36 harbored early-pseudotime STICs with minimal resistance-associated signaling, whereas patient 43 (Figure 2E) displayed transcriptionally advanced STICs enriched for DNA repair, cell cycle, and PARPi resistance programs while maintaining low EMT activity. p53 signature lesions in patients with HGSC showed low EMT activity in patients 6, 9, 10, and 14 but reached STIC-like levels in patients 17, 20 and 25, consistent with their more advanced pseudotime positions (Figure 2D, Supplementary Figure S2B). By contrast, p53 signature lesions in patients without HGSC showed predominantly low pathway activity, with the exception of patient 34, who exhibited pathway enrichment also in matched normal fimbrial tissue (Figure 2F). Finally, unsupervised k-means clustering identified subgroups of STIC phenotypes characterized by differential pathway engagement (Figure 2G, Supplementary Table-S9), further highlighting heterogeneity across STICs. STIC clustering patterns were distinct from those observed for HGSC lesions (Figure S2C, Supplementary Table-S10).

## Discussion

Using trajectory-based transcriptomic analysis, we demonstrate that STICs are not a uniform precursor state but instead occupy distinct evolutionary positions along a continuum from normal epithelium to invasive HGSC. Although STICs are widely regarded as obligate precursors of HGSC, our data demonstrate marked interpatient heterogeneity. STICi span early to later pseudotime states, frequently presenting loss of cilium organisation. By contrast, STICc consistently exhibit transcriptional programs characteristic of more advanced malignant progression, including cell cycle and mitotic programs, EMT/morphogenesis, interferon response and DNA repair pathways. These findings suggest that STICs are heterogeneous and may have different progression risks.

Biological processes related to cilia structure and function showed the strongest decrease alongside the malignant transformation pseudotime. Although loss of primary cilium is linked to carcinogenesis through Hedgehog and Wnt signaling(14), little is known about the role and significance of motile cilia loss in ovarian carcinogenesis(15-17). Our data indicate that markers of cilium assembly and function could potentially be used to evaluate early carcinogenesis in HGSC.

Importantly, we observed enrichment of PARPi resistance–associated transcriptional signatures that varied across STICs, including within the same lesion category (STICi/STICc) and across *BRCA*-status (STICc). While many STICi occupied early pseudotime states with limited therapeutic drug resistance-associated signatures (paclitaxel/ PARPi), a subset aligned with later pseudotime positions exhibited transcriptional profiles comparable to STICc, which more consistently showed activation of DNA repair and replication stress–attenuation programs. Notably, these resistance associated states were often established at the STIC stage and persisted with only modest amplification in HGSC. EMT is a process known to associate with poor survival in HGSC (18), and we observed variable activity of the EMT-associatd programs in early lesions.. In patients with concurrent cancer, transcriptional activity of EMT could be observed as early as p53 signature lesions. Heterogeneity was also evident with respect to DNA repair activity across STICs, with STICc potentially showing higher activity compared to STICi. Together, these findings raise caution against uniform PARPi-based primary medical prevention for all STICs. Not all STICs may be equally sensitive to PARPi, raising the possibility of toxicity risks, without clear benefit in biologically indolent or intrinsically resistant lesions. This highlights the need for further research in this area.

As the first trajectory based transcriptomic analysis of STICs, this study provides an evolutionary framework but is limited by its cross-sectional design, relatively small number of cases, and lack of longitudinal clinical outcomes. Our data do not preclude that STICi could be sensitive to PARPi; however, this should be interpreted with caution given the limited sample size. In particular, expanding the number of STICi samples and capturing diverse ethnic backgrounds will be essential to more robustly define the spectrum of STIC phenotypes along the malignant evolution timeline and also to better understand their relationship to PARPi sensitivity and resistance.

Future studies in larger patient cohorts integrating multi-omics, longitudinal sampling, and outcome data will be required to refine risk stratification and prognosis, and to develop therapeutic as well as preventive strategies. Forjaz et al. recently reported the first multi-omics, whole-human Fallopian tube 3D imaging study, suggesting that STIC lesions may be substantially more common than indicated by existing protocols(19). Our findings of marked heterogeneity and differential progression among STICs are also consistent with this observation. Evolutionary trajectory-informed biomarkers may improve our ability to distinguish and identify the clinically significant lesions, enabling individualized surveillance, targeted interception, and prevention strategies using biology-driven approaches.

## Author Contributions

Project conceptualisation by MS, EM, RM, FB. Data curation and formal analysis by MS and EM. Resources by RD, TK and SS. First manuscript draft by MS and EM. Manuscript supervision by RM and FB. Manuscript review and editing by RM, FB, RD, TK and SS.

## Acknowledgements

FB and EM acknowledge support from UKRI Frontier Research grant EP/X028704/1 and City of London CRUK Core Award CTRQQR-2021\100004.

## Supplementary Figure

**Supplementary Figure 1.**
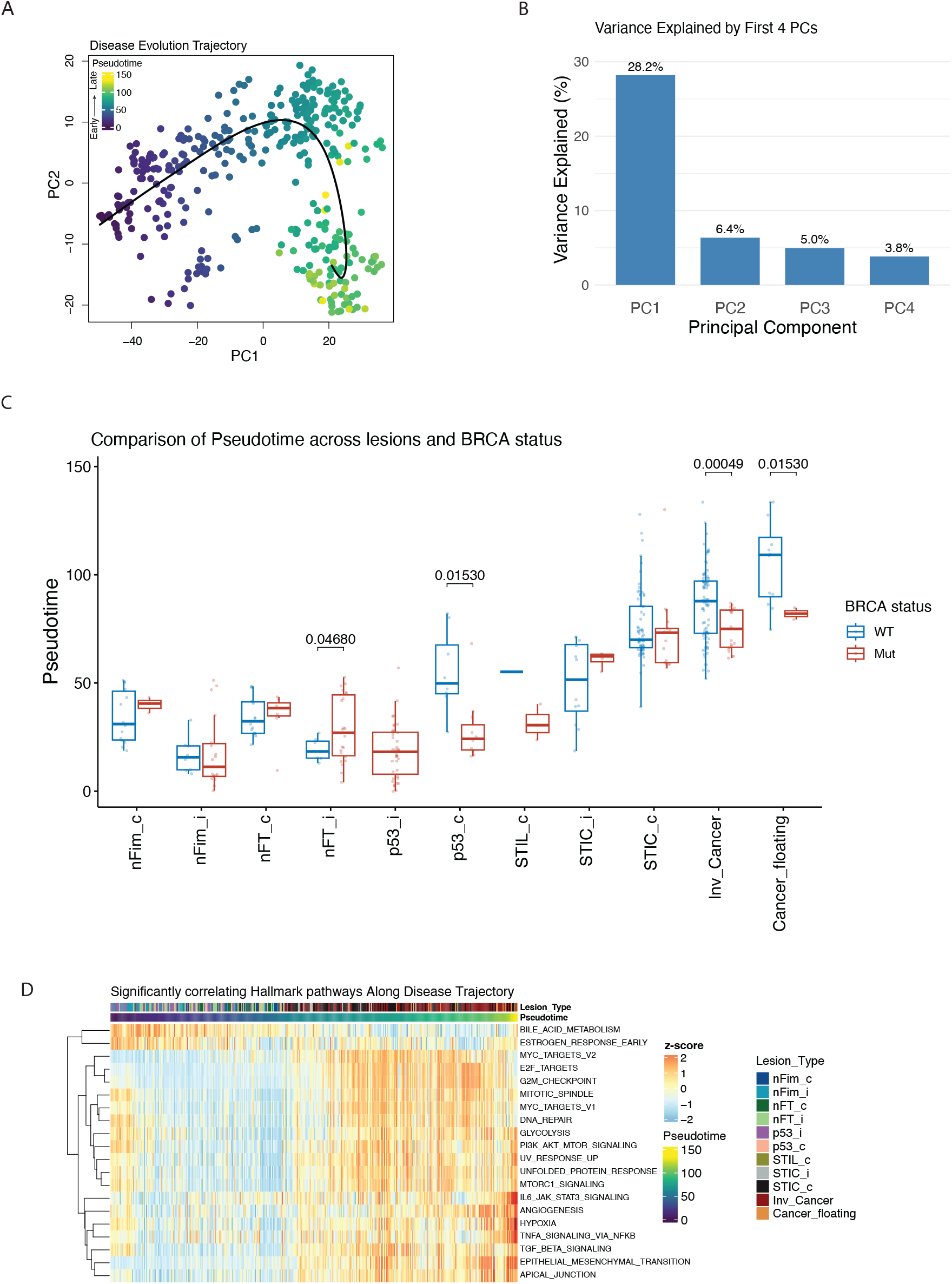
A) PCA plot of Figure 1A, coloured by the slingshot’s pseudotime value. Pseudotime corresponds to a one-dimensional variable representing each ROI’s transcriptional progression towards terminal state. **B)** Scree plot showing variance captured by the first 4 PC components used in slingshot. **C)** Boxplot illustrating ROIs pseudotime values per BRCA status for each lesion type (p values correspond to BH adjusted Welch’s t-test). **D**) Heatmap of gsva enrichment scores for Hallmark pathways that significantly change in expression with pseudotime, along the first slingshot trajectory (Spearman BH adjp < 0.05, r > 0.3)

**Supplementary Figure 2.**
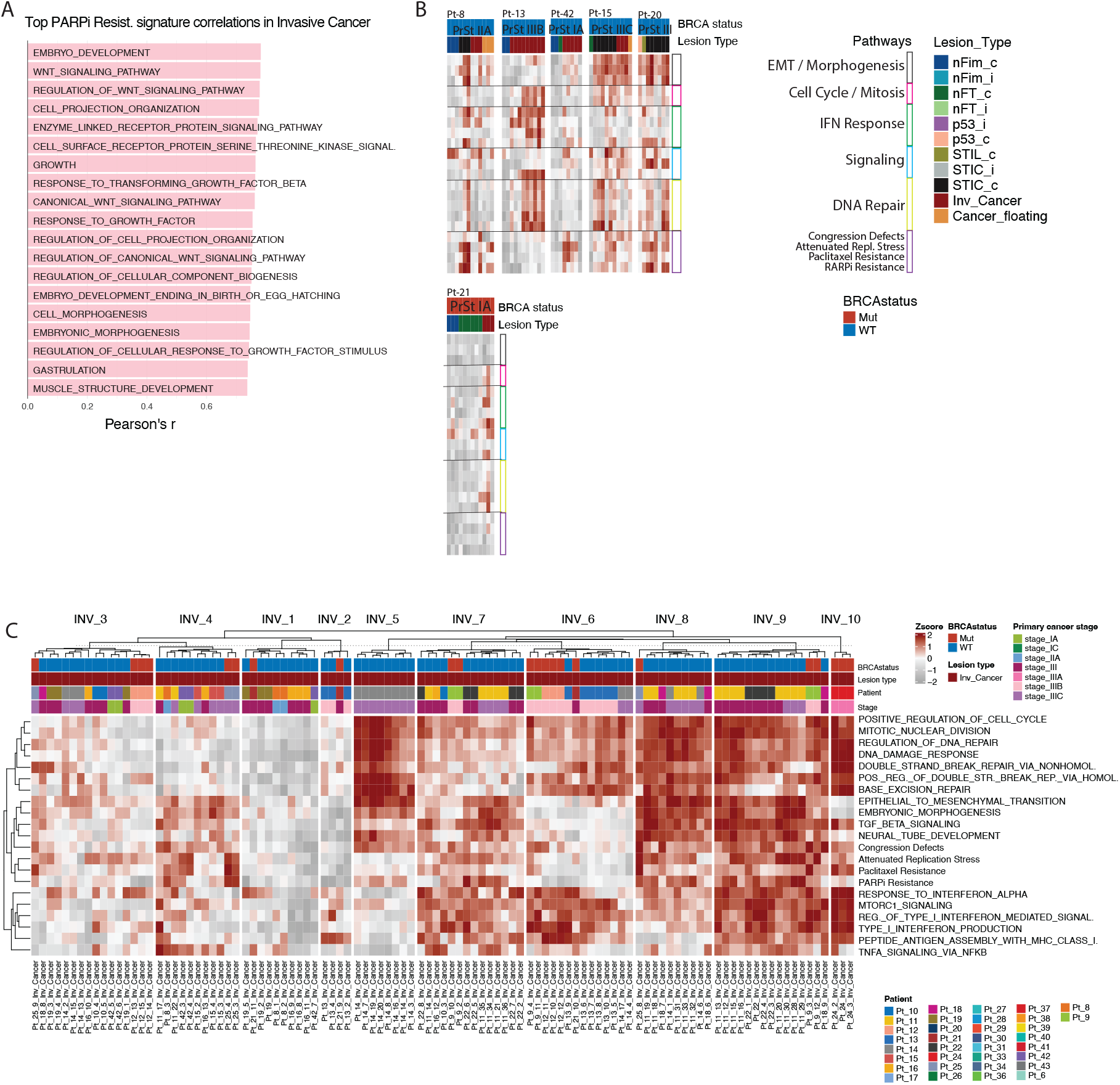
**A)** Barplot of Invasive Cancer ROI transcriptomes showing top 10 GOBPs significantly correlating with a PARP-inhibitor resistance signature from Tamura et al (pearson’s BH adjp < 0.05). **B)** Heatmaps of significant disease related GOBPs and Hallmarks as well as the Tamura et al transcriptional signatures for patients with invasive cancer. Primary cancer stage information is also noted on the heatmaps. **C)** Heatmap of gsva enrichment scores showing clustering patterns of invasive cancer ROIs using the processes and pathways described in Figure 2 panel (D-F). Clustering of ROIs was performed by k-means clustering.

